# The N-cadherin interactome in primary cardiomyocytes as defined by quantitative proximity proteomics

**DOI:** 10.1101/348953

**Authors:** Yang Li, Chelsea D. Merkel, Xuemei Zeng, Jonathon A. Heier, Pamela S. Cantrell, Mai Sun, Donna B. Stolz, Simon C. Watkins, Nathan A. Yates, Adam V. Kwiatkowski

## Abstract

The junctional complexes that couple cardiomyocytes must transmit the mechanical forces of contraction while maintaining adhesive homeostasis. The adherens junction (AJ) connects the actomyosin networks of neighboring cardiomyocytes and is required for proper heart function. Yet little is known about the molecular composition of the cardiomyocyte AJ or how it is organized to function under mechanical load. Here we define the architecture, dynamics and proteome of the cardiomyocyte AJ. Mouse neonatal cardiomyocytes assemble stable AJs along intercellular contacts with organizational and structural hallmarks similar to mature contacts. We combine quantitative mass spectrometry with proximity labeling to identify the N-cadherin (CDH2) interactome. We define over 350 proteins in this interactome, nearly 200 of which are unique to CDH2 and not part of the E-cadherin (CDH1) interactome. CDH2-specific interactors are comprised primarily of adaptor and adhesion proteins that promote junction specialization. Finally, we find evidence of dynamic interplay between AJ and Z-disc proteins. Together, our results provide novel insight into the cardiomyocyte AJ and provide a proteomic atlas for defining the molecular complexes that regulate cardiomyocyte intercellular adhesion.

**Summary Statement:** Proximity proteomics reveals a specific and specialized N-cadherin (CDH2) interactome along the cell-cell contacts of primary cardiomyocytes.

## Introduction

Heart function requires mechanical coupling and chemical communication between cardiomyocytes through a specialized adhesive structure called the intercalated disc (ICD). The ICD is formed from three junctional complexes: adherens junctions (AJs) and desmosomes that physically link opposing cardiomyocytes, and gap junctions that electrically couple cardiomyocytes (Ehler, 2016; Vermij et al., 2017; Vite and Radice, 2014). AJs and desmosomes link the actin and intermediate filament (IF) cytoskeletons, respectively, to the ICD and provide structural integrity and mechanical strength to the cell-cell contact. ICD formation requires multiple adhesion, cytoskeletal and signaling proteins, and mutations in these proteins can cause cardiomyopathies (Ehler, 2018). However, the molecular composition of ICD junctional complexes remains poorly defined.

The core of the AJ is the cadherin-catenin complex (Halbleib and Nelson, 2006; Ratheesh and Yap, 2012). Classical cadherins are single-pass transmembrane proteins with an extracellular domain that mediates calcium-dependent homotypic interactions. The adhesive properties of classical cadherins are driven by the recruitment of cytosolic catenin proteins to the cadherin tail: p120-catenin (CTNND1) binds to the juxta-membrane domain and β-catenin (CTNNB1) binds to the distal part of the tail. β-Catenin, in turn recruits αE-catenin (CTNNA1) to the cadherin-catenin complex. α-Catenin is an actin-binding protein and the primary link between the AJ and the actin cytoskeleton (Drees et al., 2005; Pokutta et al., 2014; Rimm et al., 1995; Yamada et al., 2005). In mice, loss of AJ proteins in the heart – N-cadherin (CDH2), β-catenin, αE-catenin or αT(Testes)-catenin (CTNNA3) – causes dilated cardiomyopathy (Kostetskii et al., 2005; Li et al., 2012b; Li et al., 2005; Sheikh et al., 2006). Mutations in αT-catenin, an α-catenin homolog expressed predominantly in the heart and testes, and the β-catenin homolog plakoglobin (JUP) have been linked to arrhythmogenic right ventricular cardiomyopathy (Li et al., 2012a; van Hengel et al., 2013), as have disruptions in β-catenin signaling (Garcia-Gras et al., 2006).

The AJ is best understood in the context of epithelia, where it regulates intercellular adhesion, cell motility and polarity (Garcia et al., 2018; Padmanabhan et al., 2015). The AJ can both sense and respond to mechanical force (Charras and Yap, 2018; Hoffman and Yap, 2015), though the molecular mechanism remains largely undefined. In epithelia, the AJ associates with a panoply of proteins that regulate adhesion, signaling and protein turnover. Recent proteomic studies have begun to define the cadherin interactome and have offered new insight into the molecular complexes that regulate AJ biology in epithelia (Guo et al., 2014; Van Itallie et al., 2014). Yet it is unclear if these complexes are shared between cell types or whether specific proteins are recruited to AJs to meet specific physiological needs. For example, in cardiomyocytes the AJ is thought to anchor myofibrils to the ICD to transmit force between cells. If and how the cardiomyocyte AJ proteome is tuned to meet the mechanical demands of contraction is not known.

Here we describe efforts to define the molecular complexes associated with N-cadherin at cardiomyocyte cell-cell contacts. We use a combination of light and electron microscopy to reveal that primary neonatal cardiomyocytes assemble junctional complexes along developing intercellular contacts with structural hallmarks reminiscent of mature contacts in adult tissue. We show that the AJ proteins are stable but dynamic, similar to epithelia. We use proximity proteomics to identify N-cadherin-associated proteins along cardiomyocyte cell-cell contacts. We define a robust repertoire of interactors, comprised primarily of adaptor and adhesion proteins unique to cardiomyocytes. Our results offer novel insight into the critical adhesion complexes that connect cardiomyocytes and provide a proteomic platform for deciphering how molecular complexes are organized to regulate cardiomyocyte adhesion and cellular organization.

## Results

### Organization of primary cardiomyocyte intercellular contacts

Primary cardiomyocytes isolated from rodent neonates retain the ability to establish cell-cell contacts in culture (Franke et al., 2007; Goncharova et al., 1992). Neonatal cardiomyocytes from mice are also amenable to transient transfection and adenoviral infection (Ehler et al., 2013). To begin to define the junctional complexes at these newly formed contacts, we first examined the recruitment of endogenous CDH2, the core of the AJ. Mouse neonatal cardiomyocytes were isolated from P0-P2 pups and plated on isotropic Collagen I substrates at high density to promote intercellular interactions. After 2-3 days in culture, neonatal cardiomyocytes had established CDH2-positive contacts around much of their perimeter (Fig. 1A, B). Myofibril formation is evidenced by the periodic, sarcomeric organization of the Z-line marker α-actinin (ACTN2) (Fig. 1C). Notably, CDH2 localization is not uniform along contacts; instead, it is discontinuous (Fig. 1B, asterisks) and often concentrated at sites of myofibril coupling between cells (Fig.1B, white arrows).

**Figure 1.**
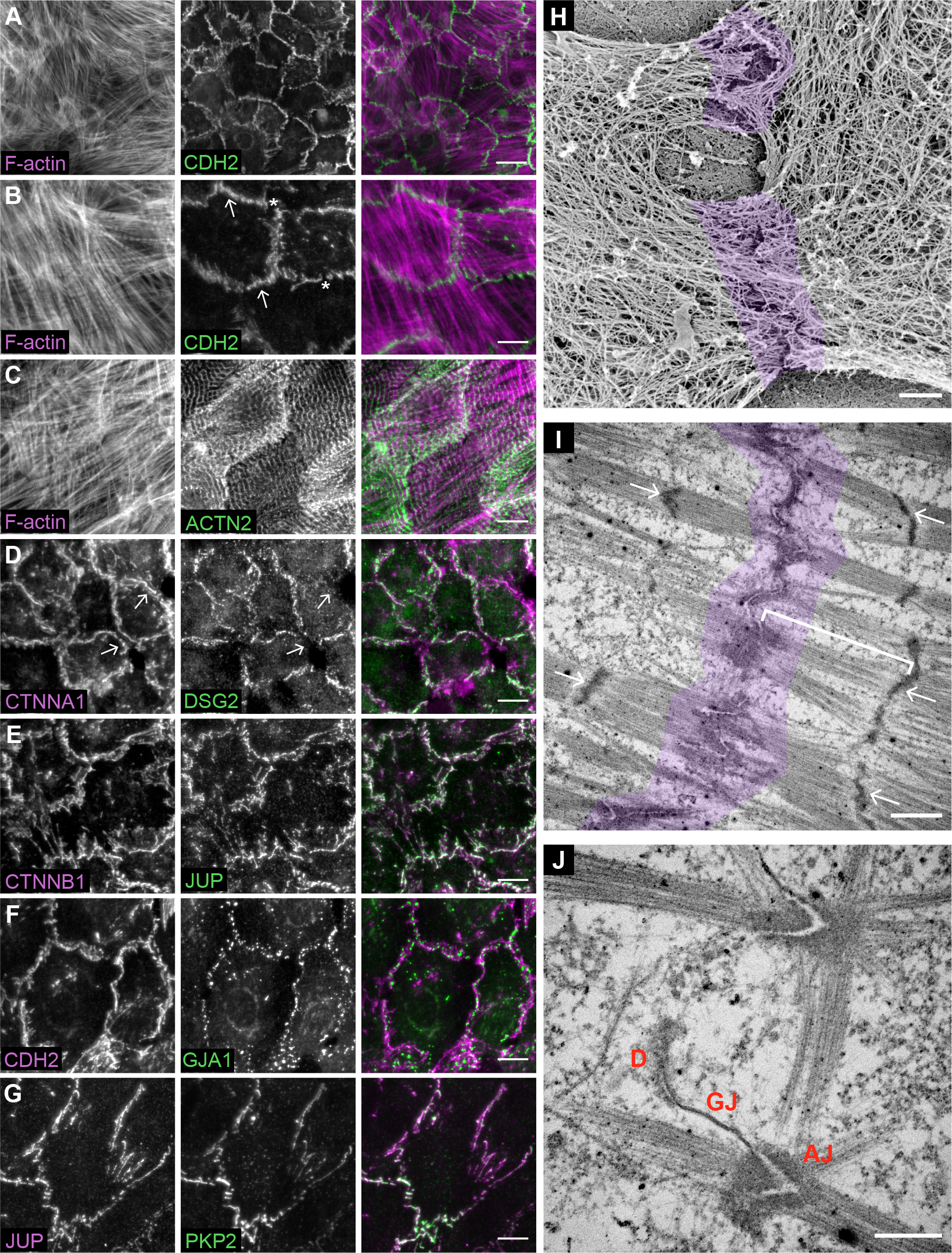
Cardiomyocyte cell-cell contact organization and architecture. A-G. Mouse neonatal cardiomyocytes plated to confluency, fixed 48-72 hrs post-plating and stained for: F-actin and CDH2 (A & B), F-actin and ACTN2 (C), CTNNA and DSG2 (D), CTNNB1 and JUP (E), CDH2 and GJA1 (F), and JUP and PKP2 (G). Individual channels and merge shown. In (B), white asterisks and white arrows mark gaps and sites of myofibril integration, respectively, along contacts. In (D), white arrows mark CTNNA1 positive, DSG2 negative contacts. H. Platinum replica electron microscopy image of two connected cardiomyocytes. Cell-cell contact is highlighted in purple. I-J. Thin section electron microscopy images of cardiomyocyte cell-cell junctions. In (I), the cell-cell contact is highlighted in purple, white arrows point to Z-discs and the white bar defines a membrane-proximal sarcomere. In (J), desmosome (D), gap junction (GJ) and adherens junction (AJ) are labeled. Scale bar is 20 μm in A; 10 μm in B-G; 500 nm in H; 1 μm in I; 500 nm in J.

We then examined the localization of other primary components of the ICD junctional complexes: AJ, desmosomes and gap junctions. As expected, the AJ proteins CTNNA1 (Fig. 1D) and CTNNB1 (Fig. 1E) showed patterns of localization identical to CDH2. The desmosomal cadherin desmoglein 2 (DSG2) also concentrated at cell-cell contacts, but its localization was more restricted than the AJ with some contacts lacking DSG2 (Fig. 1D, white arrows mark CTNNA1 positive, DSG2 negative contacts). Interestingly, two proteins typically associated with the desmosome – JUP and plakophilin 2 (PKP2) – showed patterns of localization nearly identical to the adherens junction (Fig. 1E, G). Finally, the gap junction protein Connexin 43 (GJA1) showed a punctate pattern of localization along contacts (Fig. 1F). Thus, the primary ICD junctional complexes are recruited to neonatal cardiomyocyte cell-cell contacts.

We then sought to define the actin architecture at contacts. We used platinum replica electron microscopy (PREM) to examine actin organization with single filament resolution (Svitkina, 2017). Cardiomyocytes are enshrouded by a dense cortical cytoskeleton that masks the underlying myofibril network and its association with junctional complexes (Fig. 1H, junctions highlighted in purple). We then used thin section transmission electron microscopy (TEM) to examine junction architecture. Thin section TEM revealed myofibrils coupled along electron-dense contacts (Fig. 1I, J; junction highlighted in purple). The contacts are highly convoluted, and many junctions adopt a chevron-like appearance (Fig. 1J). In addition to adherens junctions, desmosomes and gap junctions are also observed (Fig. 1J), consistent with the immunostaining. Importantly, the junctional topology of cultured cardiomyocytes is similar to that observed in adult hearts, where the angled junctions may help to balance shear versus tensile stresses during contraction (Bennett et al., 2006). Taken together, we conclude that neonatal cardiomyocytes build junctional complexes with many of the organizational and structural hallmarks of adult heart tissue.

### Adherens junction protein dynamics

We next examined the dynamics of CDH2 and associated catenin proteins in cardiomyocytes. GFP-tagged CDH2, CTNNB1, JUP, CTNNA1 and CTNNA3 were individually transfected into cardiomyocytes. All fusion constructs localized to cell-cell contacts, as expected (Fig. 2A). Protein dynamics were measured by fluorescent recovery after photobleaching (FRAP) in dense cells that had been plated for 48-72 hours (Fig. 2A). Fluorescence recovery over ten minutes was quantified, plotted and fit to double exponential curve (Fig. 2B). The mobile fractions of junctional CDH2 (34.4%), CTNNB1 (32.3%), JUP (26.5%), CTNNA1 (36.4%) and CTNNA3 (36.1%) were all similar to each other (Fig. 2C). Notably, these fractions were nearly identical to those observed in epithelial cells (Yamada et al., 2005) indicating that the majority (~2/3) of cadherin/catenin complexes are immobile components of the AJ plaque.

**Figure 2.**
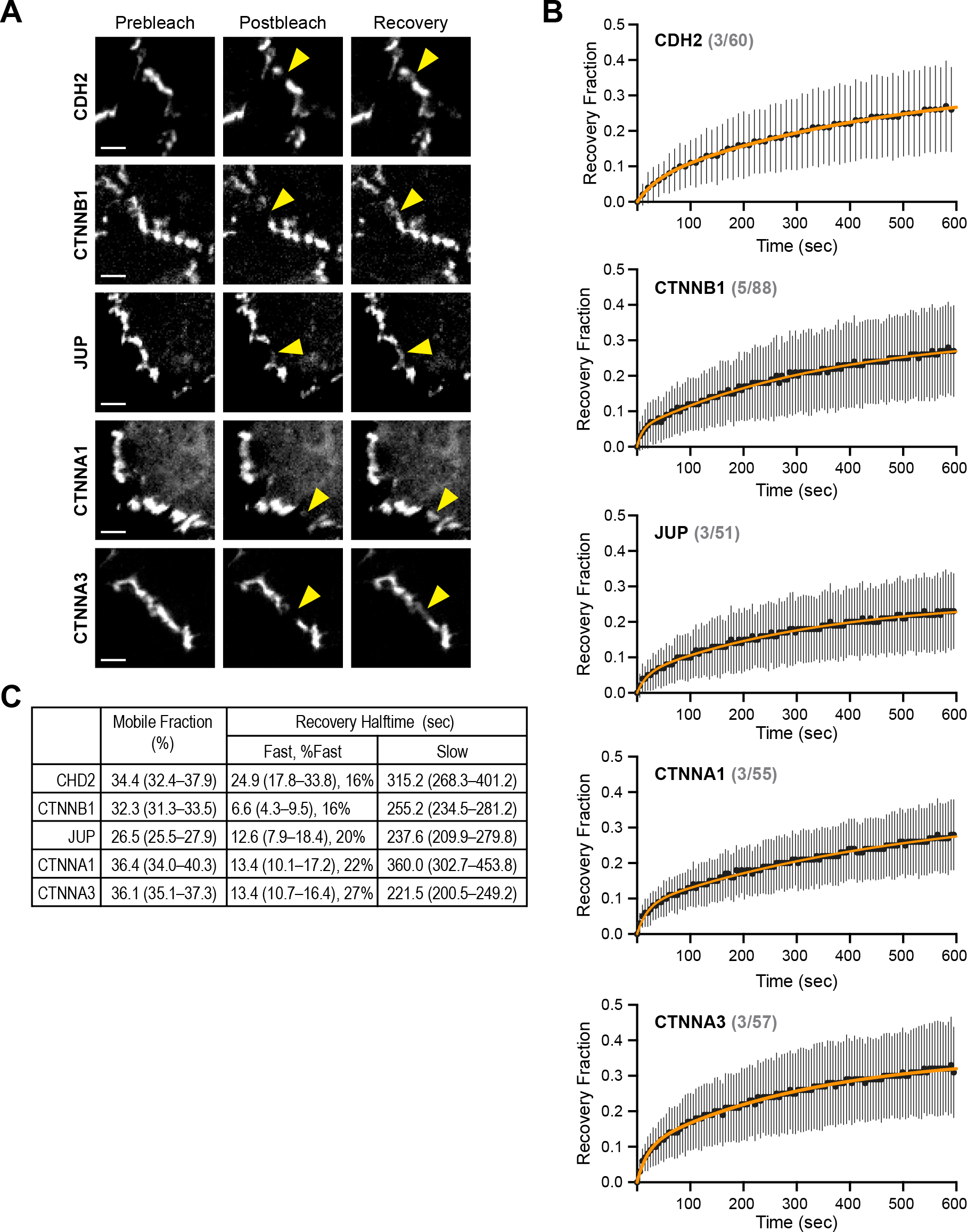
Adherens junction protein dynamics at cardiomyocyte cell-cell contacts. A. Representative prebleach, postbleach and recovery images from FRAP studies of cells expressing GFP-tagged CDH2, CTNNB1, JUP, CTNNA1 or CTNNA3. Yellow arrows mark the FRAP region along a cell contact. B. Plot of FRAP recovery fraction over time. At each time point, the mean recovery fraction is shown as a black circle and the standard deviation is represented by black lines. The data was fit to a double exponential curve (orange line). The number of experiments and number of FRAP contacts quantified for each protein is shown in grey (# experiments/# FRAP contacts) C. Summary of the mobile fraction (as percentage) and recovery halftimes (fast and slow pools). The percentage of the fast pool also listed. Scale bar is 50 μm in A.

We then assessed the recovery rates of the mobile fractions for both the fast and slow pools (Fig. 2C). For the cytoplasmic catenins, the fast pool recovery halftimes (6.6–13.4 sec) could reflect an unbound, cytosolic population of protein near cell contacts. Alternatively, the pool could be caused by photoswitching (Mueller et al., 2012). The fast pool recovery of the transmembrane CDH2 (24.9 sec) likely represents photoswitching because we do not expect diffusion of new CDH2 during this initial time frame. Importantly, the fast pool for all components is relatively small (16–27%) and the slow pool represents the dynamics of the majority of the junction population. Here, the half-times of fluorescence recovery were also similar: CDH2 (315.2 sec), CTNNB1 (255.2 sec), JUP (237.6 sec), CTNNA1 (360.0 sec) and CTNNA3 (200.5 sec). This reflects the tight associations between core components of the cadherin-catenin complex (Pokutta et al., 2014) and suggest that the multiprotein complex is exchanged as a unit along contacts. While E-cadherin (CDH1), CTNNB1 and CTNNA1 were found to have similar rates of recovery at epithelial cell-cell contacts, the rates were approximately an order of magnitude faster, in the realm of 26–40 sec (Yamada et al., 2005). It is unclear what underlies this difference, but it could reflect differences in CDH2-mediated trans interactions (Katsamba et al., 2009; Vendome et al., 2014) and/or junctional membrane topology and its potential impact on protein diffusion. Together, our results suggest that cardiomyocytes form stable AJs with properties similar to epithelia.

### CDH2-BioID2 biotinylates proteins at cardiomyocyte cell-cell contacts

Given the unique structural and mechanical qualities of cardiomyocyte cell-cell contacts, we next sought to define the molecular complexes along the junctional membrane. We used proximity proteomics to identify proteins near CDH2 by fusing the biotin ligase BioID2 (Kim et al., 2016) to the C-terminal tail of Cdh2 (Fig. 3A). This technique has been used with success to define the CDH1 interactome in epithelia (Guo et al., 2014; Van Itallie et al., 2014) and define CTNNA1 force-dependent molecular interactions (Ueda et al., 2015). We cloned the Cdh2-BioID2 fusion into an adenoviral expression system and made Cdh2-BioID2 adenovirus that would allow us to infect primary cardiomyocytes and express low levels of Cdh2-BioID2 for imaging and protein analysis (Fig. 3B). We were able to reproducibly infect >90% of cardiomyocytes at a low multiplicity of infection (MOI). The Cdh2-BioID2 fusion localized to cell-cell contacts (HA stain, Fig. 3C), similar to endogenous CDH2 (Fig. 1A, B). Importantly, when biotin (50 μM) was added to the culture, Cdh2-BioID2 labeled proteins along cell-cell contacts (SA stain in Fig. 3E; compare to uninfected control in Fig. 3D). Biotin addition and concomitant labeling did not disrupt cell-cell contacts (Fig. 3E) and optimal biotinylation was achieved after 24 hours (Fig. S1). In addition to the prominent junction labeling, a smaller population of biotinylated proteins was observed at Z-discs (Fig. 3F, G). Finally, we were able to precipitate biotinylated proteins from lysates of infected cells cultured with biotin (Fig. 3H). Thus, Cdh2-BioID2 localizes to cardiomyocyte cell-cell contacts and labels proximal proteins that can be isolated for proteomic analysis.

**Figure 3.**
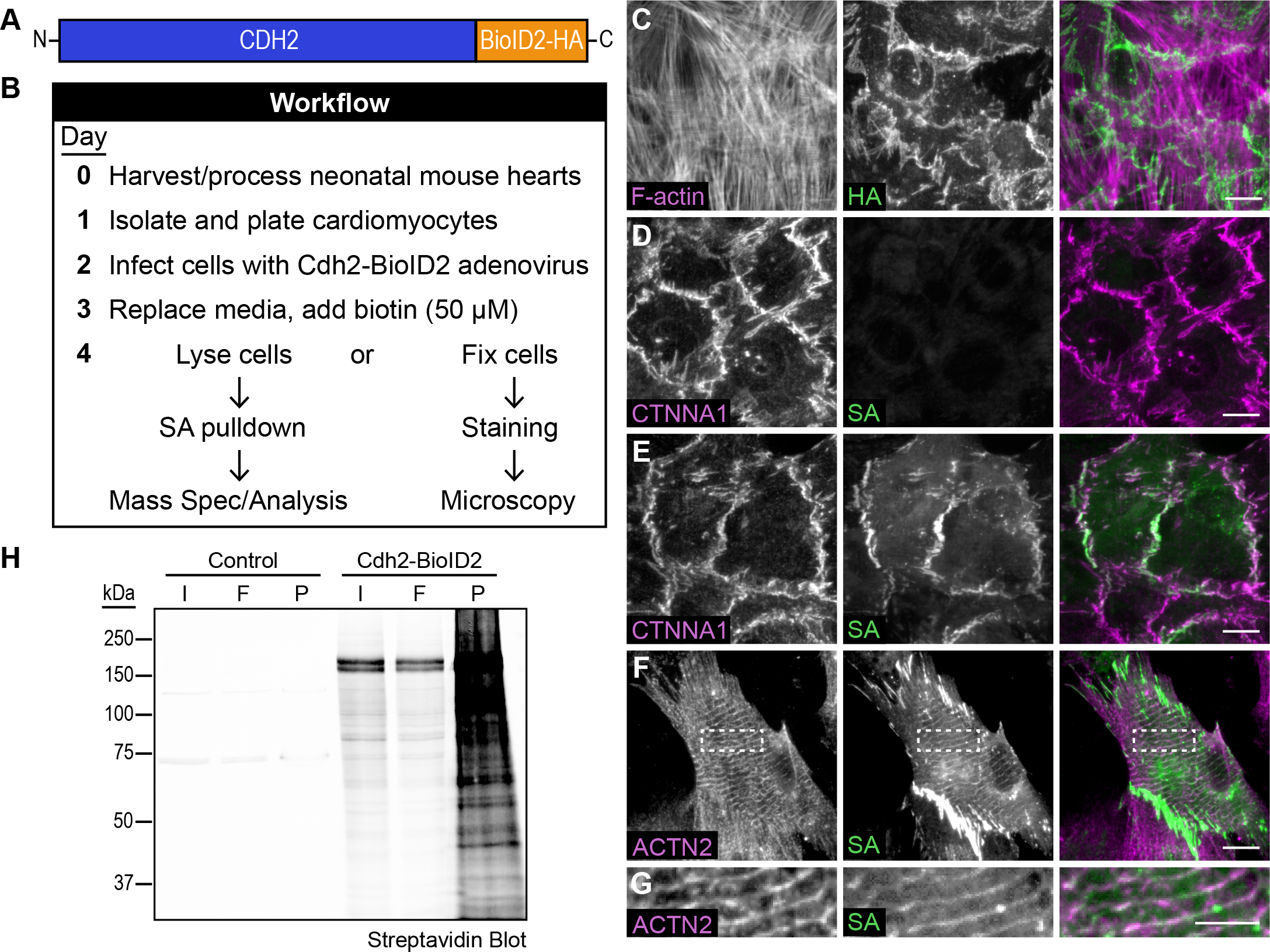
CDH2-BioID2 localizes to cell contacts and labels junctional proteins. A. Cartoon schematic of CDH2-BioID2 fusion. B. Experimental workflow for infecting primary cardiomyocytes, labeling with biotin and protein fixation/isolation. C-G. Staining of Cdh2-BioID2 infected cardiomyocytes. C. Cdh2-BioID2 infected cardiomyocytes were stained for F-actin (magenta in merge) and HA (green in merge) to identify the HA-tagged fusion construct. D, E. Uninfected (D) and Cdh2-BioID2 infected (E) cardiomyocytes were stained for CTNNA1 and labeled with a streptavidin (SA) conjugated to CY3 to identify biotinylated proteins. F, G. Cdh2-BioID2 infected cardiomyocytes stained for ACTN2 and biotin (SA). (G) is a high mag image of the boxed region in (F) highlighting biotinylated proteins along Z-lines. H. Streptavidin western blot of pulldowns from control and Cdh2-BioID2 infected cardiomyocytes. Initial material (I), flow through (F) and precipitated material (P) marked. Scale bar is 10 μm C-F; 5 μm in G.

### Quantitative proximity proteomics reveals the cardiomyocyte CDH2 interactome

We used quantitative mass spectrometry (MS) to define the CDH2 interactome. For each replicate, 4 million cells were infected with Cdh2-BioID2 adenovirus, biotin was added and the cells were harvested following the workflow in Fig. 3B. Uninfected control samples were treated identically to Cdh2-BioID samples (i.e., 50 μM biotin added 48 hrs post-plating and cells harvested 24 hrs after biotin addition). Six Cdh2-BioID2 replicates and six control replicates were collected and analyzed.

MS sample analysis revealed a total of 5117 peptides from 917 proteins (Fig. 4A, B). The mean coefficient of variance for the Cdh2-BioID2 replicates was ~30% (Fig. S2). When single unique peptides were excluded, the list was reduced to 4687 peptides from 487 proteins (Fig. 4B). To define Cdh2-BioiD2 enriched proteins, we established thresholds of fold change ≥ 10 and p < 0.001 (Fig. 4A, dashed grey lines). These thresholds culled the list to a final 365 proteins from 354 genes (Fig. 4B; Table S1).

**Figure 4.**
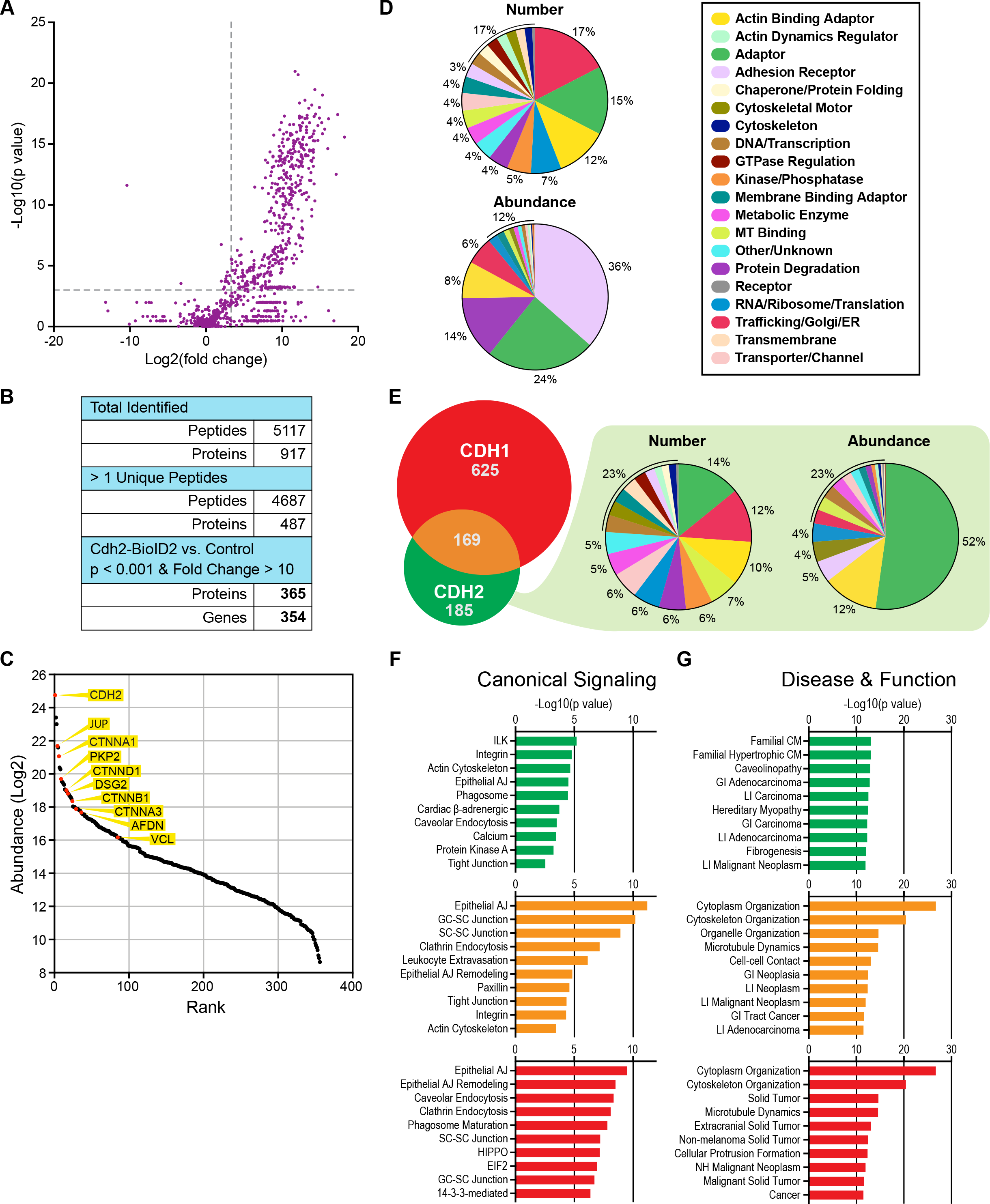
Quantitative mass spectrometry identifies CDH2 interactome. A. Plot of p value (−Log_10_) versus fold change (Log_2_) of identified proteins. Dashed grey lines marks p = 0.001 (y axis) and fold change = 10 (x axis). B. Summary of identified proteins. C. Rank plot of abundance (iBAQ mass, Log_2_). Proteins of interest are marked as red circles and labeled. D. Protein distribution by assigned category based on number (top pie chart) or abundance (iBAQ) (bottom pie chart). E. Venn diagram of CDH2 interactome in cardiomyocytes (green) versus CDH1 interactome from epithelial cells (red). 169 proteins are shared (orange). Distribution of the CDH2 only pool (minus CDH2, 184 proteins) based on number (left) or abundance (right). F, G. IPA enrichment analysis of CDH2 only (green), CDH2/CDH1 shared (orange) and CDH1 only (red) groups in canonical signaling pathways (F) or disease and function (G). Abbreviations: AJ, Adherens Junction; CM, Cardiomyopathy; GC, Germ Cell; GI, Gastrointestinal; LI, Large Intestine; NH; Nonhematologic; SC, Sertoli Cell.

The relative abundance of these 365 proteins is plotted in Fig. 4C and the 35 most abundant proteins are listed in Table 1. Among the most abundant proteins were core components of the AJ, including CTNNB1, JUP, CTNND1 and CTNNA1. These same proteins were also abundant in the CDH1 interactome (Guo et al., 2014; Van Itallie et al., 2014). Notably, the desmosome components DSG2 and PKP2 were also abundant hits, as were CTNNA3 and the α-catenin ligands vinculin (VCL) and afadin (AFDN) (Hazan et al., 1997; Pokutta et al., 2002; Tachibana et al., 2000; Weiss et al., 1998). We speculate that the abundance of desmosomal proteins DSG2 and PKP2 could reflect the proximity of AJs and desmosomes along developing cardiomyocyte junctions and/or the proposed intermingling of junctional components in hybrid junctions (Franke et al., 2006). The enrichment of VCL and AFDN, two actin-binding proteins that help anchor the AJ to actin (Sawyer et al., 2009; Yonemura et al., 2010), likely reflects the importance of these proteins in connecting the AJ to the myofibril network.

**Table 1.** 35 most abundant proteins in the CDH2 interactome.

| Name | Description | Category | p-Value | Fold change | Mass Relative to CDH2 |
| --- | --- | --- | --- | --- | --- |
| CDH2 | Cadherin 2, N-cadherin | Adhesion receptor | 4.8E-10 | 487 | 1.000 |
| RPS27A | Ribosomal protein S27a | Protein degradation | 4.2E-12 | 168 | 0.393 |
| SORBS2 | Sorbin and SH3 domain containing 2 | Adaptor | 2.4E-06 | 2578 | 0.159 |
| JUP | Junction plakoglobin | Adaptor | 7.1E-07 | 63 | 0.120 |
| AHNKA | AHNAK nucleoprotein | Adaptor | 5.8E-10 | 1454 | 0.112 |
| CTNNA1 | Catenin alpha-1, αE-catenin | Actin binding adaptor | 2.3E-10 | 4243 | 0.078 |
| PICALM | Phosphatidylinositol binding clathrin assembly protein | Trafficking/Golgi/ER | 6.1E-06 | 2858 | 0.048 |
| RTN4 | Reticulon-4 | Trafficking/Golgi/ER | 1.2E-09 | 1171 | 0.045 |
| PKP2 | Plakophilin 2 | Adaptor | 3.2E-05 | 959 | 0.030 |
| NEBL | Nebulette | Actin binding adaptor | 2.7E-06 | 3905 | 0.027 |
| XIRP1 | Xin actin binding repeat containing 1 | Actin binding adaptor | 5.9E-05 | 380 | 0.026 |
| ITGB5 | Integrin subunit beta 5 | Adhesion receptor | 2.7E-16 | 295308 | 0.025 |
| CSRP3 | Cysteine and glycine rich protein 3 | Actin binding adaptor | 3.5E-04 | 222 | 0.024 |
| ANXA2 | Annexin A2 | Membrane binding adaptor | 5.5E-06 | 24 | 0.019 |
| CTNND1 | Catenin delta 1, p120 catenin | Adaptor | 6.5E-06 | 996 | 0.019 |
| BICD2 | BICD cargo adaptor 2 | MT binding | 4.0E-05 | 909 | 0.018 |
| AHNAK2 | AHNAK nucleoprotein 2 | Adaptor | 2.9E-09 | 1014 | 0.017 |
| DSG2 | Desmoglein 2 | Adhesion receptor | 2.3E-07 | 5522 | 0.016 |
| NACA | Nascent polypeptide-associated complex $\alpha$ subunit | RNA/Ribosome/Translation | 1.3E-06 | 3137 | 0.016 |
| MYH6 | Myosin 6 | Cytoskeletal motor | 3.2E-06 | 11 | 0.015 |
| MAP4 | Microtubule associated protein 4 | MT binding | 6.7E-07 | 6048 | 0.015 |
| EEF1A1 | Eukaryotic translation elongation factor 1 alpha 1 | RNA/Ribosome/Translation | 6.4E-08 | 24 | 0.014 |
| PLIN4 | Perilipin 4 | Metabolic Enzyme | 1.2E-08 | 394 | 0.013 |
| CTNNB1 | Catenin beta 1, β-catenin | Adaptor | 3.0E-13 | 136626 | 0.012 |
| SNAP23 | Synaptosome associated protein 23 | Trafficking/Golgi/ER | 4.0E-07 | 1336 | 0.010 |
| CRIP2 | Cysteine rich protein 2 | DNA/Transcription | 5.1E-07 | 1103 | 0.009 |
| PALM2 | Paralemmin 2 | Adaptor | 2.9E-06 | 1414 | 0.009 |
| SHROOM1 | Shroom family member 1 | Actin binding adaptor | 1.8E-05 | 1022 | 0.009 |
| CTNNA3 | Catenin alpha 3, αT-catenin | Actin binding adaptor | 5.4E-08 | 6617 | 0.009 |
| CAVIN1 | Caveolae associated protein 1 | Trafficking/Golgi/ER | 8.3E-07 | 1093 | 0.009 |
| LRRC59 | Leucine rich repeat containing 59 | RNA/Ribosome/Translation | 4.6E-06 | 720 | 0.009 |
| EFHD2 | EF-hand domain family member D2 | Adaptor | 3.4E-06 | 868 | 0.008 |
| FHL1 | Four and a half LIM domains 1 | Adaptor | 1.8E-04 | 319 | 0.008 |
| MYL3 | Myosin light chain 3 | Cytoskeletal motor | 4.5E-04 | 38 | 0.008 |
| PAKAP | Paralemmin A kinase anchor protein | Adaptor | 3.3E-05 | 354 | 0.008 |

### The cardiomyocyte CDH2 interactome is distinct from the epithelial CDH1 interactome

We assigned each of the 354 genes in the Cdh2-BioID2 interactome into one of 20 functional categories according to information from Uniprot, GeneCards and Entrez (Fig. 4D), similar to (Guo et al., 2014). By number, the categories with the most hits were Trafficking/Golgi/ER (17%), Adaptor (15%) and Actin Binding Adaptor (12%). However, when considering protein abundance (iBAQ), the top categories were Adhesion Receptor (36%), Adaptor (24%), Protein Degradation (14%) and Actin Binding Adaptor (8%) (Fig. 4D). Given the substantial, electron-dense structures built along cardiomyocyte AJs (Fig. 1I, J), the abundance of adaptor proteins and adhesion receptors could function to couple myofibrils between cardiomyocytes.

We then compared the Cdh2-BioID2 hits with CDH1 interactome from epithelia (Guo et al., 2014; Van Itallie et al., 2014). There are 169 proteins shared between the two interactomes (Fig. 4E) and 185 proteins unique to CDH2 in cardiomyocytes. The distribution of the CDH2-only hits was similar to the entire population (Fig. 4D), with adaptor proteins forming the largest class in number and abundance (Fig. 4E). Actin-binding adaptors, adhesion receptors and cytoskeletal motor proteins were also enriched in the CDH2-only pool (Fig. 4E). By abundance, adaptor proteins account for 64% of the CDH2-only pool, highlighting the specialized molecular machinery required for intercellular adhesion in cardiomyocytes.

To gain further insight into the potential similarities and differences between CDH2 and CDH1 interactomes, we performed enrichment analysis using Ingenuity Pathway Analysis (IPA). We examined the CDH2, CDH1 and CDH2/CDH1 protein sets in canonical signaling and disease & function pathways. The CDH1 and CDH2/CDH1 sets were both enriched for AJ, cell-cell and endocytosis signaling (Fig. 4F; Table S2). In contrast, the CDH2-specific pool showed less enrichment overall, though the emergence of cardiac β-adrenergic and calcium signaling pathways could reflect how the CDH2 interactome is tuned to cardiac function (Fig. 4F; Table S2). The top enriched disease & function pathways for the CDH1 and CDH2/CDH1 protein sets were cellular organization and cancer-related categories (Fig. 4G; Table S3). In contrast, the CDH2-specific pool was enriched for a variety of cardiomyopathies (Fig. 4G; Table S3). These results suggest that, in cardiomyocytes, CDH2 recruits and organizes unique molecular complexes to regulate cell-cell adhesion and signaling.

### Differential gene expression contributes to the specialized adhesion complexes in cardiomyocytes

During development, differential gene expression plays an essential role in establishing cell identity and function. Underlying their specialized role in cardiac contraction, cardiomyocytes express a unique set of genes. To determine if differential gene expression contributes to the CDH2 interactome, we identified cardiomyocyte or heart enriched genes in gene expression profiling data and compared these enriched genes to the CDH2, CDH1 and shared protein sets. We identified 1319 cardiomyocyte-enriched genes (CEGs) from RNA sequencing (RNA-seq) data collected at 11 points during the differentiation of human induced pluripotent stem cells (hiPSCs) to cardiomyocytes (Tompkins et al., 2016). Comparative analysis revealed that 52 CEGs were unique to CDH2 (Fig. 5A), representing 28.1% (52/185) of the CDH2 hits (Fig. 5C). In contrast, the number of CEGs present in the CDH1 or CDH2/CDH1 sets was much lower, representing just 4.6% and 15.3%, respectively, of the hits for each class (Fig. 5C). We also calculated the percentage of CDH2, CDH1 and CDH2/CDH1 CEGS in the total CEG pool (1319 CEGs). Fisher’s exact test indicated that CEGs were highly enriched in the CDH2 set, but not the CDH1 or CDH2/CDH1 sets (Fig. 5C).

**Figure 5.**
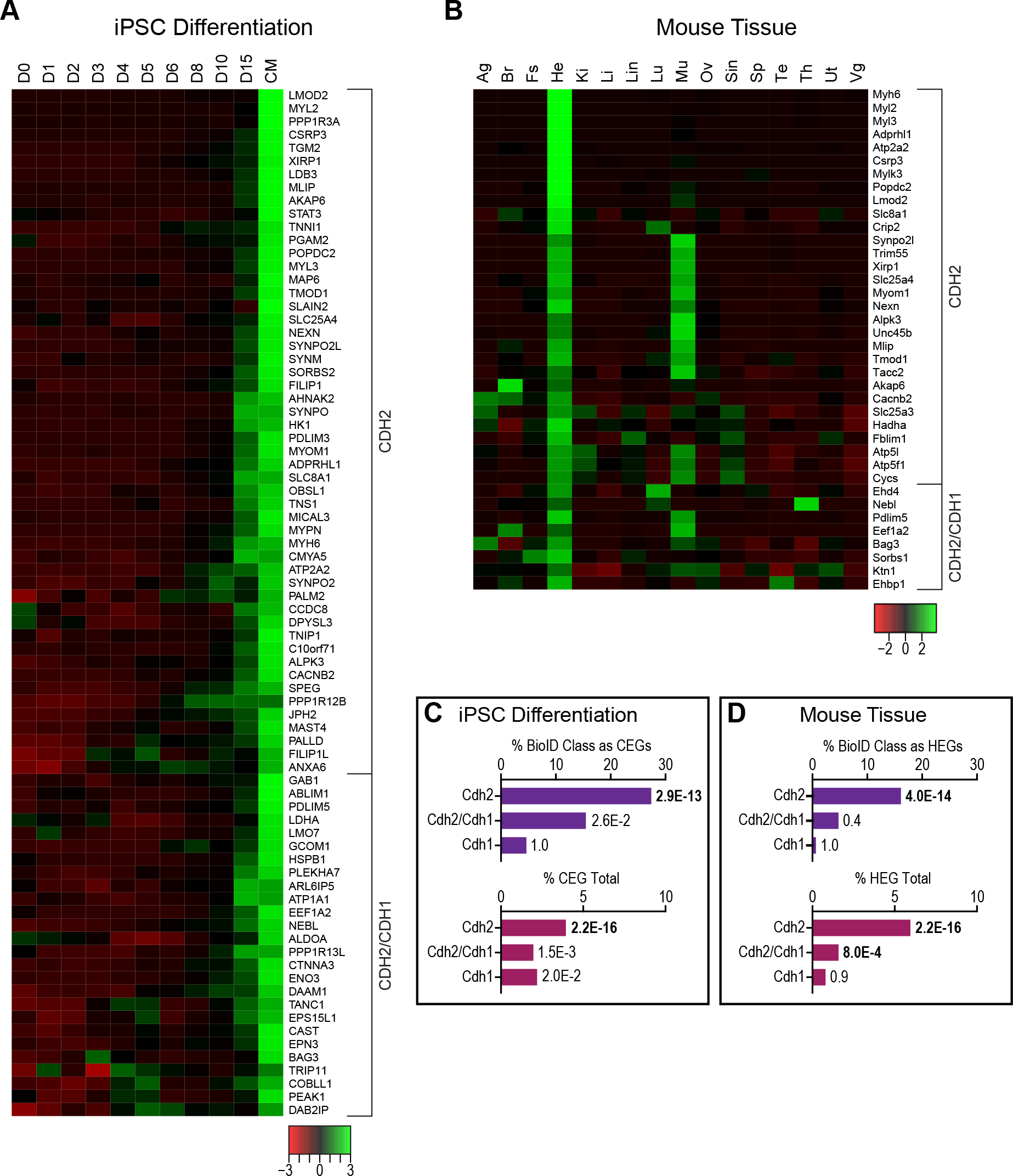
Differential gene expression contributes to the cardiomyocyte CDH2 proteome. A. Heat map of CDH2 or CDH2/CDH1 expression profiles during iPSC differentiation into cardiomyocytes (CM), D0 (day 0) to D15 (day 15). B. Heat map of CDH2 or CDH2/CDH1 expression profiles in mouse tissues. Ag (adrenal gland), Br (brain), Fs (fore stomach), He (heart), Ki (kidney), Li (liver), Lin (large intestine), Lu (lung), Mu (muscle), Ov (ovary), Sin (small intestine), Sp (spleen), Te (testis), Th (thymus), Ut (uterus) and Vg (vesicular gland). C, D. Top, % BioID Class as CEGS/HEGs: percentage of each BioID class as cardiomyocyte enriched genes (CEGs) or heart enriched genes (HEGs). Bottom, % CEG Total: fraction of those BioID CEGs/HEGs in the total CEG/HEG population. P value of Fisher’s exact test shown. Significant values are in bold.

We also analyzed tissue-enriched genes from adult mice. Using RNA-seq data from mouse tissues (Li et al., 2017), we identified 504 heart-enriched genes (HEGs). Of those, 42 HEGs were present in the CDH2 set, representing 22.7% of the CDH2 hits and 8.3% of total HEGs (Fig. 5B, D). No HEG enrichment was observed in the CDH1 or CDH2/CDH1 sets. Similar results were observed in gene expression data from human tissues (Fig. S3). Together, these results suggest that cardiomyocyte and heart signature gene expression contribute significantly to the CDH2 interactome.

### CDH2 interactome protein network

To better understand how molecular complexes could be assembled at cardiomyocyte AJs, and how these complexes might differ from epithelia AJs, we connected and organized the CDH2 interactome (Fig. 6A). We defined a new interactome group – ICD proteins (curated from the human protein atlas (Estigoy et al., 2009)) – and compared it to the CDH2 and CDH1 interactomes. The three-way comparison (Fig. 6B) defined four groups of proteins: CDH2, CDH2/CDH1, CDH2/ICD and CDH2/CDH1/ICD (Fig. 6A). All proteins were assigned a symbol based on primary function and color-coded to match their group. We then constructed a hierarchical classification with CDH2 at the top (see Methods for details). All protein-protein interactions were based on published, experimental data. The classification produced four tiers of interactors: 11 primary, 62 secondary, 177 tertiary and 48 quaternary (Fig. 6A). 52 of the Cdh2-BioID hits could not be connected to any other protein in the network (Table S4). The hierarchal organization reveals that the percentage of Cdh2-BioID unique hits (green) increases from 0 to 70% as the distance from CDH2 increases, whereas the percentage of CDH2/CDH1 (orange) and CDH2/CDH1/ICD (pink) groups decreases from >90% to 25% (Fig. 6C, D). This suggests that the primary complex (1° and 2° tiers) is largely shared between CDH2 and CDH1, but that specific, specialized interactors are recruited outside (3° and 4° tiers) the primary complex to regulate junction assembly in cardiomyocytes. Also noteworthy is the abundance of CDH2 (green) hits versus CDH2/ICD (purple) or CDH2/CDH1/ICD (pink) hits. These green-labeled proteins reflect potentially new, previously unassigned ICD components with potential roles in cadherin and cardiomyocyte adhesion biology.

**Figure 6.**
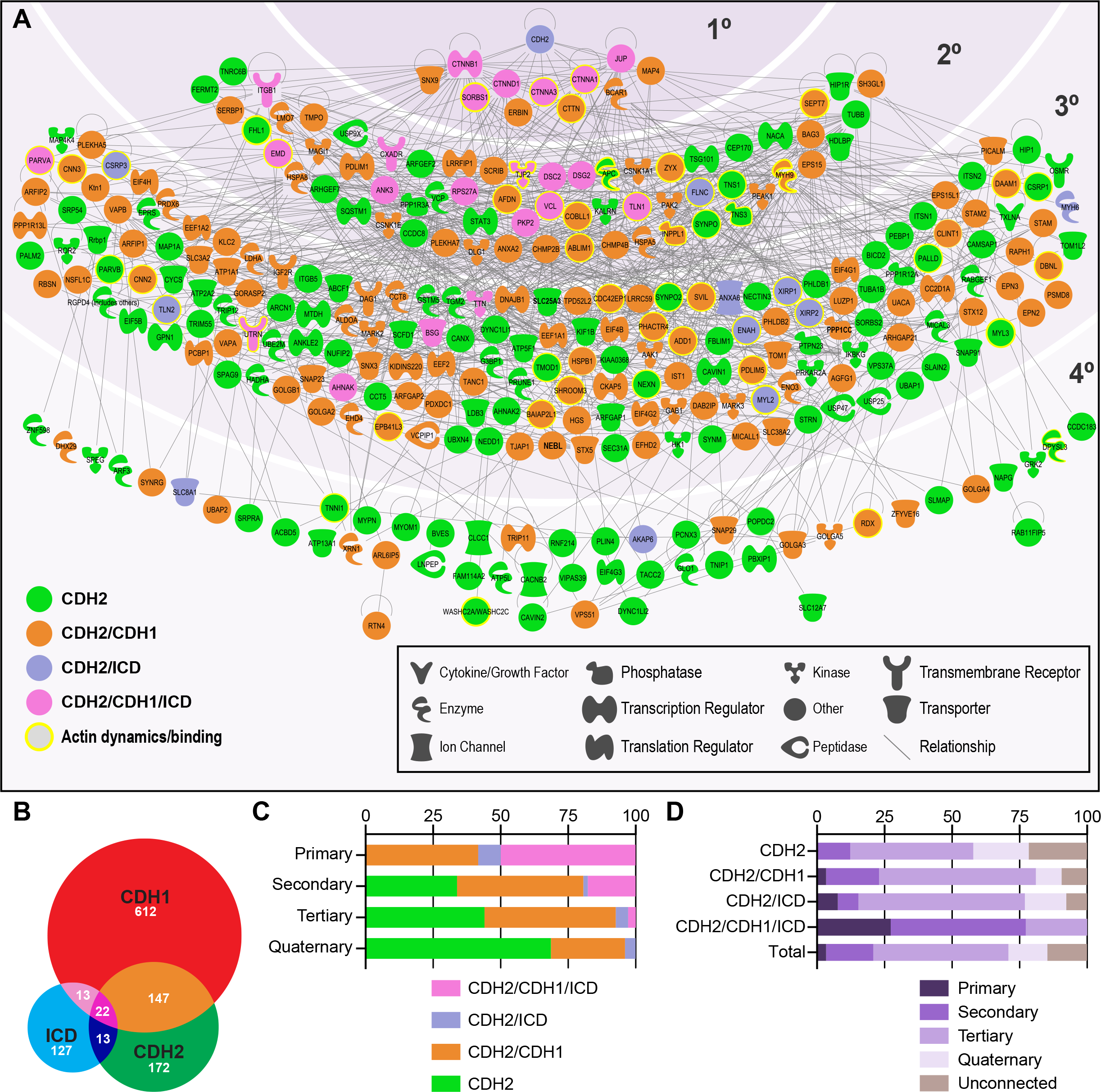
Cardiomyocyte CDH2 interactome. A. Interaction network of CDH2 interactome organized into four tiers based on IPA protein-protein interactions with experimental evidence. Bottom left legend defines group classification (Venn diagram in B); bottom right legend defines symbol shape and protein function. B. Venn diagram between CDH2 interactome, CDH1 interactome and ICD curated proteins. C. Distribution of groups within each interactome tier. D. Distribution of tier and unconnected proteins within each group and the total collection.

### Identified actin-binding proteins localize to cell-cell contacts and Z-discs

The primary function of the AJ is to link the actin cytoskeletons of neighboring cells. In cardiomyocytes, the AJ must connect contractile myofibrils, placing unique demands on the proteins that physically connect actin to the cadherin complex. We identified nearly 50 actin-binding or actin-associated proteins in the CDH2 interactome, spread across multiple tiers (Fig. 6A, actin-related proteins highlighted with yellow; Table S1) including the actin-binding proteins VCL and AFDN, top-ranked hits (Fig. 4C and Table 1). We examined the localization of a number of these actin-binding proteins by transiently expressing GFP-tagged versions in cardiomyocytes (Fig. 7). Zonula Occludens 1 (TJP1; note that paralog TJP2 was identified in this screen), Dishevelled Associated Activator of Morphogenesis 1 (DAAM1), and Cortactin (CTTN) all localized primarily to cell-cell contacts (Fig. 7A-C). AFDN localized primarily to cell-cell contacts but was also present at Z-discs (Fig. 7D). VCL localized to cell-cell contacts as well as cell-substrate contacts (Fig. 7E). Supervillin (SVIL), Synaptopodin 2 (SYNPO2) and Emerin (EMD) all localized primarily to Z-discs, but they were also observed colocalizing with CDH2 at contacts (Fig. 7F-H; Fig. S4).

**Figure 7.**
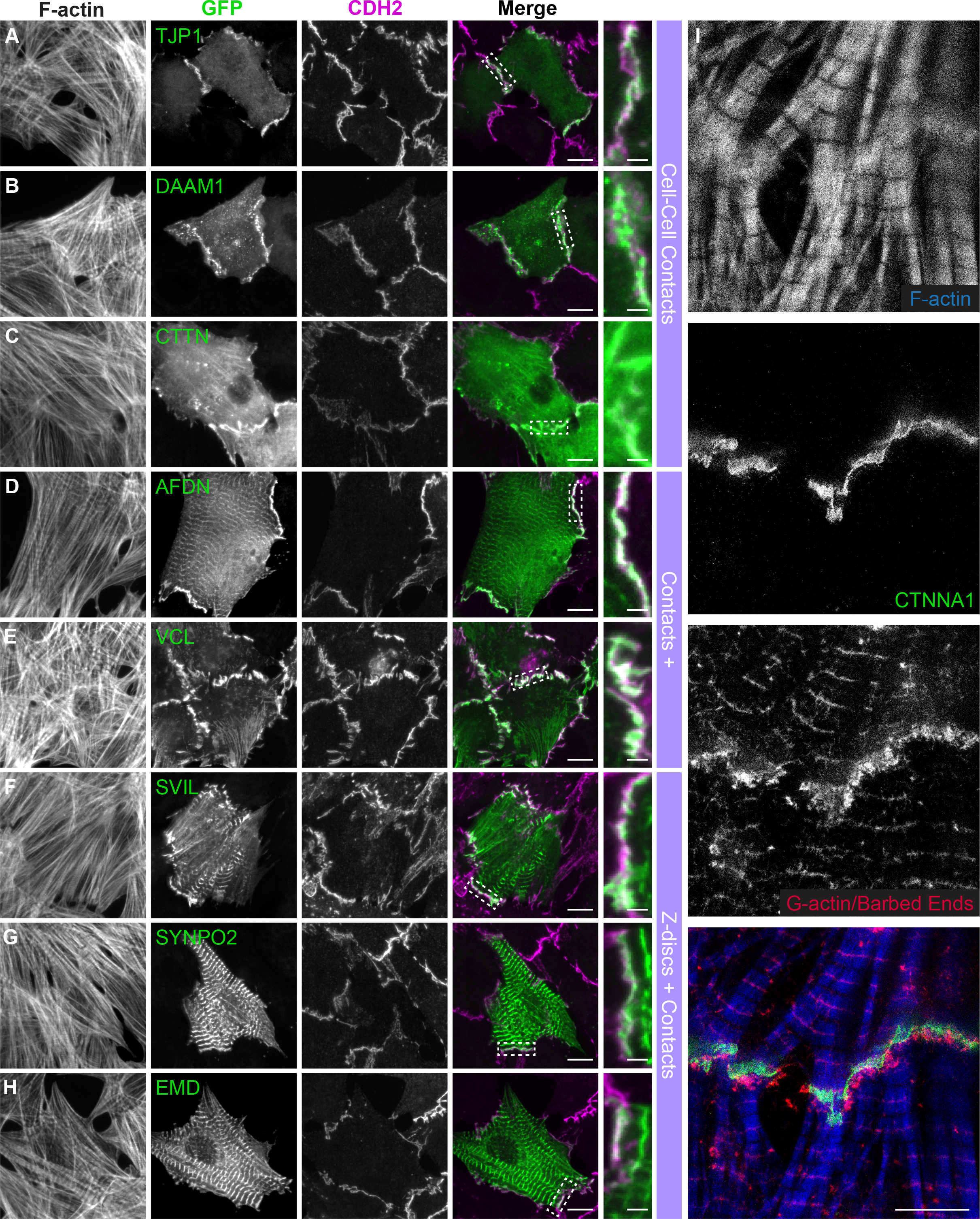
Identified actin-binding proteins localize to cell-cell contacts and z-discs. A-H. Cardiomyocytes transfected with GFP-tagged Cdh2-BioID hits. Cells were fixed 24 hours post-transfection and stained for Cdh2 and F-actin. Individual and merged GFP (green) and CDH2 (magenta) channels shown. Far right column is a magnification of boxed contact in Merge image. I. STED image of cardiomyocyte with F-actin barbed ends labeled and stained for F-actin and CDH2. Scale bar is 10 μm in Merge in A-H; 2 μm in magnified view in A-H; 2 μm in I.

In Cdh2-BioID2 expressing cardiomyocytes, biotinylated proteins were detected at cell-cell contacts as well as at Z-discs (Fig. 1 3G, H). The ICD is sometimes referred to as the terminal Z-disc because, for a membrane-tethered myofibril, the ICD functions as the terminal end of the sarcomere (Fig. 1I). We performed a barbed end assay to analyze the polarity of actin filaments in cardiomyocytes. We permeabilized cardiomyocytes in the presence of fluorescent G-actin, fixed and then stained for F-actin and CTNNA1. We observed prominent labeling at Z-discs and, notably, along cell-cell contacts (Fig. 7I). We speculate that Z-disc proteins are recruited dynamically to the AJ to promote myofibril assembly. Thus, the ICD AJ plays an important role in guiding both cardiomyocyte adhesion and cytoskeleton organization.

## Discussion

Our results provide new details into the architecture of the developing ICD and define the proteins that organize the AJ in cardiomyocytes. This work builds off past studies of the cadherin-catenin interactome in epithelia (Guo et al., 2014; Ueda et al., 2015; Van Itallie et al., 2014) to expand the AJ protein atlas to include the cardiomyocyte, a unique contractile system. Our molecular and proteomics data reveal how ancillary proteins associate with the cadherin-catenin complex to build the cardiomyocyte AJ and guide cardiomyocyte organization and adhesion.

### Core adhesion complexes are conserved

The components and dynamics of the core AJ complex in cardiomyocytes are similar to epithelia. The cadherin-catenin complex is recruited to developing contacts and its components are among the most robust hits in the proteomic screen (Table 1). Likewise, FRAP studies revealed that the complex is largely immobile (~1/3 is mobile) and that the entire complex is exchanged along contacts similar to epithelia (Yamada et al., 2005), though the rate of the turnover is ~10 fold slower. These properties are not unexpected given current AJ dogma. Biochemical studies have demonstrated that β-catenin binds with high affinity to αE-catenin and that the β-catenin/ αE-catenin complex binds strongly to the cadherin tail to create a stable complex (Pokutta et al., 2014). These molecular interactions are apparently conserved in cardiomyocytes, despite the unique contact architecture and physical demands of muscle contraction. We propose that cardiomyocyte AJ protein behavior is dictated primarily by the basic principles of cadherin and catenin biology.

The desmosome components DSG2, JUP and PKP2 were among the more abundant proteins isolated in the proteomic screen (Table 1). DSG2 localization was more restricted than the AJ and appeared to be preferentially localized near sites of myofibril integration (Fig. 1D; Merkel and Kwiatkowski, unpublished observations). DSG2 (and desmosome development) may be favored at more stable or mature AJs, consistent with EM analysis (Fig. 1J). In contrast, JUP and PKP2 showed near identical localization patterns to the AJ (Fig. 1E, G). JUP can bind directly to classical cadherins (Choi et al., 2009) and was a robust hit in CDH1 proteomic screens (Guo et al., 2014; Van Itallie et al., 2014). PKP2, an armadillo protein related to CTNND1, is a multifunctional protein that binds desmosomal cadherins, JUP and desmoplakin (DSP) (Bass-Zubek et al., 2009). We speculate that both JUP and PKP2 may be recruited to AJs during the initial stages of contact formation to promote desmosome assembly, similar to their proposed role in epithelia (Nekrasova and Green, 2013). Note that PKP2 can also bind directly to CTNNA3 and has been proposed to link the AJ to intermediate filaments at hybrid junctions, where AJ and desmosome proteins are mixed in mammalian hearts (Borrmann et al., 2006; Goossens et al., 2007).

### AJ specialization is driven by ancillary adaptor proteins

We identified 365 proteins from 354 genes in the CDH2 interactome. Of these, 169 hits were shared with the CDH1 interactome while 185 hits were unique to CDH2. These CDH2-specific hits were enriched in genes linked to cardiac signaling and cardiomyopathies, reflective of the specialization of the CDH2 interactome to meet the demands of cardiomyocyte physiology. By abundance, 60% of the CDH2 pool was composed of adaptor proteins. Analysis of the protein-protein interactions reveals that the many shared components occupy the inner tiers of the network. In contrast, CDH2-specifc adaptors primarily occupy the outer tiers of the protein network. This suggests that the cardiomyocyte AJ recruits a unique set of cytoskeletal, scaffold and signaling proteins to build this critical mechanical junction and regulate adhesive homeostasis in a demanding contractile system.

We found that the CEGs or HEGs contribute to approximately 20-30% of the differential components in cardiomyocyte AJs. The remaining 70-80% of unique components in cardiomyocyte AJs reflects differential organization at the protein level. Thus, while differential gene expression is a significant contributor to interactome identity, the primary driver of AJ specialization is the recruitment of adaptor proteins to build specific, multiplex protein complexes.

Roughly 140 curated ICD proteins were not detected in the CDH2 interactome (Fig. 7B), including established CDH2-associated proteins like the gap junction protein Connexin 43 (GJA1). While we observed GJA1 along contacts in our cultured cardiomyocytes, the localization was punctate and differed from CDH2. The absence of GJA1 and other curated ICD proteins from our screen could be due to the physical limitations of BioID2-labeling (the range of biotinylation is limited to ~10 nm, (Kim et al., 2016)), the absence of surface lysines on a target protein for labeling and/or the maturity of the cell-cell contacts in our system. Alternatively, it could reflect the segregation of molecular complexes along these contacts and highlight the specificity of AJ interactions. Nonetheless, of the 185 CDH2-specific hits, only 13 were curated ICD proteins and the remaining 172 hits were not previously associated with the ICD or the AJ, thus expanding the atlas of proteins associated with cell adhesion in cardiomyocytes.

The assembled interaction network (Fig. 6A) reflects the hierarchy of protein binding required to form the molecular complexes along cardiomyocyte contacts. Critical to these assemblages are the catenin proteins, CTNND1, CTNNB1, JUP, CTNNA1 and CTNNA3. All catenin proteins are known to bind directly to a number of other proteins. For example, in addition to binding CTNNB1/JUP and actin, CTNNA1 can interact with VCL, AFDN, PKP2 and ZO1/2 (Vite et al., 2015). These proteins, in turn, can associate with a wide-range of cytoskeletal and signaling proteins. We speculate that the catenins coordinate the organization of molecular complexes at the cardiomyocyte AJ to regulate adhesion and signaling.

### The AJ and the Z disc, linked through the myofibril sarcomere

Cardiomyocytes must be coupled to create a functional syncytium and the AJ serves as the mechanical link between myofibrils and the ICD membrane. The contractile unit of the myofibril is the sarcomere whose lateral boundaries are defined by Z-discs where the barbed ends of actin filaments are interdigitated and crosslinked. Z-discs are connected to the lateral membrane (sarcolemma) and the surrounding extracellular matrix by specialized adhesive complexes called costameres. In cardiomyocytes, the AJ functions as the terminal Z-disc for the membrane proximal sarcomere (Fig. 1I). While cardiomyocyte organization almost certainly requires coordination between the AJ and Z-disc/costamere (Bennett et al., 2006), the molecular details remain largely unexplored and undefined. CDH2 proximity labeling revealed decoration of both cell-cell contacts and Z-discs (Fig. 3F, G), indicating that proteins were being biotinylated at AJs and then shuttling to Z-discs over time. Consistent with this, Z-disc/costamere proteins such as SVIL (Oh et al., 2003) and SYNPO2 (Weins et al., 2001) were identified in the proteomic screen and localized at both Z-discs and contacts (Fig. 7F, G; Fig. S4). EMD is best associated with the nuclear membrane and ICD AJ (Barton et al., 2015; Cartegni et al., 1997), but also displayed prominent Z-disc localization when expressed in cardiomyocytes (Fig. 7H). Additional proteins with Z-disc localization were identified in the CDH2 interactome including: BAG3 (Hishiya et al., 2010), LDB3 (Faulkner et al., 1999), NEBL (Moncman and Wang, 1995), PDLIM3 & PDLIM5 (Zheng et al., 2010), FHL1 (Frank et al., 2006), TTN (Herzog, 2018) and ZYX (Frank and Frey, 2011)(Table S1). Likewise, the integrins ITGB1 and ITGB5 and talins TLN1 and TLN2, core components of costamere (Jaka et al., 2015; Samarel, 2005), were identified in this screen. Interestingly, BAG3, NEBL, PDLIM5 and TTN were identified in the CDH1 interactome (Guo et al., 2014; Van Itallie et al., 2014), suggesting that these associations are conserved across cell types. We speculate that the AJ recruits Z-disc proteins to coordinate myofibril assembly and integration at contacts. Additional studies are expected to reveal how such coordination is regulated at the molecular level.

### The developing ICD in neonatal cardiomyocytes

We took advantage of the innate ability of primary neonatal cardiomyocytes to reestablish cell-cell contacts *in situ* to express tagged AJ proteins and explore their dynamics and label CDH2-associated proteins. Primary neonatal cardiomyocytes plated on isotropic substrates form cell-cell contacts around their entire perimeter (Fig. 1), similar to cardiomyocytes in the developing and perinatal heart (Hirschy et al., 2006). The stereotypical, elongated cardiomyocyte morphology with aligned myofibrils and ICDs restricted to the bipolar ends develops postnatally (Angst et al., 1997; Hirschy et al., 2006; Pieperhoff and Franke, 2007), though the mechanisms of this polarization remain unclear. Thus, while our proteomic results offer a snapshot of the CDH2 interactome at developing cell-cell contacts rather than at mature ICDs, this transitional stage has *in vivo* relevance and these results provide a significant advance in defining the cadherin interactome in cardiomyocytes. In addition, we were able to generate quantitative MS data from a relatively small sample of cultured primary cardiomyocytes. A similar experimental protocol could be used to examine changes in the CDH2 interactome from mutant cardiomyocytes or from cardiomyocytes cultured under varying conditions (e.g., soft versus stiff substrates). Alternatively, it could be used to define the CDH2 proteome in differentiated iPSCs or modified to express in an AAV system to examine the CDH2 proteome *in vivo* during heart development or disease. Future work is expected to build on this newly defined AJ network to provide important insight into how the molecular complexes that regulate AJ function change in response to injury or disease.

## Materials and Methods

### Plasmids

Murine *Cdh2* in pEGFP-N1 (CDH2-EGFP) was a gift from James Nelson. CTNNA3-EGFP was described previously (Wickline et al., 2016). Plasmid pEGFP-C1-rat-l-afadin (AFDN) was gift from Yoshimi Takai (Nakata et al., 2007). Plasmids mEmerald-JUP-N-14 (Addgene 54133), mEmerald-beta-catenin-20 (CTNNB1, Addgene 54017), mEmerald-alpha1-catenin-C-18 (CTNNA1, Addgene 53982), mEmerald-ZO1-C-14 (TJP1, Addgene 54316) and mEmerald-Vinculin-23 (VCL, #54302) were gifts from Michael Davidson. Emerin pEGFP-C1 (EMD, Addgene 61993) was a gift from Eric Schirmer (Zuleger et al., 2011). GFP-cortactin (CTNN1, Addgene 26722) was a gift from Anna Huttenlocher (Perrin et al., 2006). EGFP-supervillin (SVIL, Addgene 13040) was gift from Elizabeth Luna (Wulfkuhle et al., 1999). MCS-BioID2-HA (Addgene 74224) was a gift from Kyle Roux (Kim et al., 2016). pAdTrack-CMV (Addgene 16405) was a gift from Bert Vogelstein (He et al., 1998).

To create the SYNPO2 and DAAM1 constructs, RNA was first isolated and purified from adult mouse heart using an RNeasy Fibrous Tissue Mini kit (Qiagen) and reverse transcribed to create cDNA using Transcriptor High Fidelity cDNA Synthesis Kit (Roche). Gene specific primers were designed against the 5’ and 3’ ends of each gene to generate full-length clones by PCR. PCR products were cloned directly into pEGFP-N1 (SYNPO2) or pEGFP-C1 (DAAM1) to create GFP fusions. Assembled clones were verified by sequencing.

### Antibodies

Primary antibodies used for immunostaining were: anti-N-cadherin (1:250, Thermo Fisher Scientific 33-3900), anti-α-Actinin (1:250, Sigma A7811), anti-Desmoglein 2 (1:250, Abcam ab150372), anti-αE-catenin (1:100, Enzo Life Science ALX-804-101-C100), anti-β-Catenin (1:100, BD Transduction Laboratories 610154), anti-γ-Catenin (1:100; Cell Signaling 2309), anti-Connexin-43 (1:100, Proteintech 15386-1-AP), anti-Plakophilin 2 (1:10, Progen 651101) and anti-HA (1:100, Sigma 11867423001). Streptavidin-Cy3 (1:300, Jackson Immunoresearch 016-160-084) was used to label biotinylated proteins. Secondary antibodies used were goat anti-mouse or anti-rabbit IgG labeled with Alexa Fluor 488, 568 or 647 dyes (1:250, Thermo Fisher Scientific). F-actin was visualized using Alexa Fluor dye conjugated phalloidin (1:100-1:250, Thermo Fisher Scientific).

### Cardiomyocyte isolation and culture

All animal work was approved by the University of Pittsburgh Division of Laboratory Animal Resources. Primary cardiomyocytes were isolated from Swiss Webster or Black 6 mouse neonates (P1-P3) as described (Ehler et al., 2013; Wickline et al., 2016). For protein isolation, Swiss Webster-derived cardiomyocytes were plated onto 35 mm dishes (1 × 10^6 cells/dish) coated with Collagen Type I (Millipore). For immunostaining, cardiomyocytes were plated onto 35 mm MatTek dishes with 10 mm insets coated with Collage Type I. Cardiomyocytes were plated in plating media: 65% high glucose DMEM (Thermo Fisher Scientific), 19% M-199 (Thermo Fisher Scientific), 10% horse serum (Thermo Fisher Scientific), 5% FBS (Atlanta Biologicals) and 1% penicillin-streptomycin (Thermo Fisher Scientific). Media was replaced 16 hours after plating with maintenance media: 78% high glucose DMEM, 17% M-199, 4% horse serum, 1% penicillin-streptomyocin, 1 μM AraC (Sigma) and 1 μM Isoproternol (Sigma). Cells were cultured in maintenance media for 2-4 days until lysis or fixation.

### Immunostaining and confocal microscopy

Cells were fixed in 4% EM grade paraformaldehyde in PHEM buffer (60 mM PIPES pH 7.0, 25 mM HEPES pH 7.0, 10 mM EGTA, pH 8.0, 2 mM MgCl_2_ and 0.12 M Sucrose) or PHM (no EGTA) buffer for 10 minutes, washed twice with PBS and then stored at 4°C until staining. Cells were permeabilized with 0.2% Triton X-100 in PBS for 4 minutes and washed twice with PBS. Cells were then blocked for 1 hour at room temperature in PBS + 10% BSA (Sigma), washed 2X in PBS, incubated with primary antibodies in PBS + 1% BSA for 1 hour at room temperature, washed 2X in PBS, incubated with secondary antibodies in PBS + 1% for 1 hour at room temperature, washed 2X in PBS and then mounted in Prolong Diamond (Thermo Fisher Scientific). All samples were cured at least 24 hours before imaging.

Cells were imaged on a Nikon Eclipse Ti inverted microscope outfitted with a Prairie swept field confocal scanner, Agilent monolithic laser launch and Andor iXon3 camera using NIS-Elements imaging software. Maximum projections of 1-2 um image stacks were created for presentation.

### Barbed end assay and STED microscopy

Cardiomyocytes were cultured on collagen-coated MatTek dishes as described. Barbed ends were labeled as described previously (Balsamo et al., 2016; Furman et al., 2007). Briefly, cells were incubated in 0.5 μM Alexa Flour 488 labeled G-actin in 20 mM HEPES pH 7.5, 138 mM KCl, 4 mM MgCl2, 3 mM EGTA, 0.2 mg/ml saponin and 10 mg/ml BSA for 5 minutes at room temperature to permeabilize and label. Cells were then fixed in PHEM buffer + 4% paraformaldehyde for 10 minutes and then stained as described above.

Stimulated emission depletion (STED) microscopy samples were fixed and stained similar to standard confocal microscopy samples. Cells were imaged on a Leica TCS SP8 STED microscope and deconvolved post-acquisition.

### FRAP experiments

FRAP experiments were conducted on a Nikon swept field confocal microscope (describe above) outfitted with a Tokai Hit cell incubator and Bruker miniscanner. Actively contracting cells were maintained at 37°C in a humidified, 5% CO_2_ environment. User-defined regions along cell-cell contacts were bleached with a 488 laser and recovery images collected every 5 or 10 seconds for 10 minutes. FRAP data was quantified in ImageJ (NIH) and average recovery plots were measured in Excel (Microsoft). FRAP recovery plots represent data from >50 contacts from at least three separate transfections of unique cell preps. All curves were fit to a double exponential formula, which fit the recovery data the best, to determine the half time of recovery and the plateau in Prism (Graphpad).

### Electron microscopy

Cardiomyocytes were cultured on collagen-coated MatTek dishes and fixed as described above. After fixation and washing, cells were incubated with 1% OsO4 for one hour. After several PBS washes, dishes were dehydrated through a graded series of 30% to 100% ethanol, and then infiltrated for 1 hour in Polybed 812 epoxy resin (Polysciences). After several changes of 100% resin over 24 hours, cells were embedded in inverted Beem capsules, cured at 37C overnight, and then hardened for 2 days at 65C. Blocks were removed from the glass dish via freeze/thaw method by alternating liquid Nitrogen and 100°C water. Ultrathin (60nm) sections were collected on to 200-mesh copper grids, stained with 2% uranyl acetate in 50% methanol for 10 minutes and 1% lead citrate for 7 minutes. Samples were photographed with a JEOL JEM 1400 PLUS transmission electron microscope at 80kV with a Hamamatsu ORCA-HR side mount camera.

For platinum replica electron microscopy (PREM), cardiomyocytes were first washed with PBS and then extracted for 3 minutes in PHEM buffer (without fixative) plus 1% TritonX-100 and 10 μM phalloidin (unlabeled). Following extraction, cells were washed 3x in PHEM buffer (without fixative) plus 5 μM phalloidin and fixed for 20 minutes in PHEM buffer plus 2% glutaraldehyde. Cells were washed and stored in PBS at 4°C until processing. Fixed samples were processed for PREM as described (Svitkina and Borisy, 1998). Replicas were imaged in grids on the JEOL JEM 1400 PLUS transmission electron microscope described above.

### Adenovirus Production

Mouse *Cdh2* ORF was first amplified from CDH2-EGFP by PCR and cloned into MCS-BioID2-HA to fuse BioID2 to the C-terminal tail of N-cadherin. The Cdh2-BioID2 fusion was then subcloned into pAdTrack-CMV plasmid. Recombinant adenovirus was produced by transforming the pAdTrack-CMV-Cdh2-BioID2 plasmid into pAdEasier-1 E.coli cells (a gift from Bert Vogelstein, Addgene 16399) (He et al., 1998). Virus packaging and amplification were performed according to the protocol described by Luo and colleagues (Luo et al., 2007). Virus particles were purified using Vivapure AdenoPACK 20 Adenovirus (Ad5) purification & concentration kit (Sartorius). Adeno-X qPCR Titration Kit (Clontech) was used to calculate virus titer by quantitative PCR on an Applied Biosystems 7900HT.

### Adenovirus infection and biotin labeling

Each experimental replicate included four 35 mm dishes with 1 × 10^6 cells each (4 × 10^6 total). Cardiomyocytes were infected one day after plating with Cdh2-BioID2 adenovirus at an MOI of 2. 24 hours later (48 hours post-plating), the media was replaced with fresh maintenance media plus 50 μM biotin in both Cdh2-BioID2 infected and control uninfected samples. The next day (72 hours post-plating), cells were harvested for protein isolation and mass spec. Cell lysate preparation and affinity purification were performed according to published protocols (Kim et al., 2016; Le Sage et al., 2016).

### Western blotting

Protein samples were separated on an 8% SDS-PAGE gel and transferred onto a PVDF membrane (Bio-Rad). The membrane was blocked in TBST + 5% BSA, washed in TBST, incubated with IRDye 680RD Streptavidin (1:1000, LI-COR) in TBST, washed twice in TBST and washed once in PBS. The membrane was scanned using a LI-COR Odyssey Infrared Imager.

### Mass spectrometry and statistical analysis

All protein samples were run on precast Mini-PROTEAN TGX 10% SDS-PAGE gels (Bio-Rad) at 120 volts for 5 min so that the proteins migrated into the gel about 1 cm^2^. Gels were stained in Coomassie blue and a single, ~1 cm gel slice was excised for each sample and submitted for processing. Excised bands were digested with trypsin as previously described (Braganza et al., 2017). Briefly, the excised gel bands were destained with 25mM ammonium bicarbonate in 50% acetonitrile (ACN) until no visual stain remained and the gel pieces were dehydrated with 100% ACN. Disulfide bonds were reduced in 10mM dithiothreitol (DTT, Sigma-Aldrich Corporation) at 56°C for 1 hour and alkylated with 55mM iodoacetamide (IAA, Sigma-Aldrich Corporation) for 45 mins at room temperature in the dark. Excess DTT and IAA were removed by dehydration in 100% ACN before rehydration in 20 ng/μL trypsin (Sequencing Grade Modified, Promega Corporation) in 25 mM ammonium bicarbonate and digested overnight at 37°C. The peptides were extracted from gel pieces in a solution containing 70% ACN/5% formic and desalted with Pierce C_18_ Spin columns (Thermo Fisher Scientific) according to manufacturer’s protocol, dried in a vacuum centrifuge and resuspended in 18 μL of 0.1% formic acid. A pooled instrument control (PIC) sample was prepared by combining 4 μL from each of the 12 samples and used to monitor instrument reproducibility.

Tryptic peptides were analyzed by nLC-MS/MS using a nanoACQUITY (Waters Corporation) online coupled with an Orbitrap Velos Pro hybrid ion trap mass spectrometer (Thermo Fisher Scientific). For each nLC-MS/MS analysis, 1 μL of peptides was injected onto a C_18_ column PicoChip 25 cm column packed with Reprosil C_18_ 3 μm 120 Å chromatography with a 75 μm ID and 15 μm tip (New Objective). Peptides were eluted off to the mass spectrometer with a 66 minute linear gradient of 2-35% ACN/0.1% formic acid at a flow rate of 300 nL/min. The full scan MS spectra were collected over mass range m/z 375-1800 in positive ion mode with an FTMS resolution setting of 60,000 at m/z 400 and AGC target 1,000,000 ms. The top 13 most intense ions were sequentially isolated for collision-induced dissociation (CID) tandem mass spectrometry (MS/MS) in the ion trap with ITMS AGC target 5,000 ms. Dynamic exclusion (90s) was enabled to minimize the redundant selection of peptides previously selected for MS/MS fragmentation.

The nLC-MS/MS data were analyzed with MaxQuant software (Cox and Mann, 2008; Tyanova et al., 2016), version 1.6.0.1. Briefly, the proteomic features were quantified by high resolution full MS intensities after retention alignment and the corresponding MS/MS spectra were searched with Andromeda search engine against the Uniprot mouse database (release November 2017, 82,555 entries) (UniProt, 2011). The mass tolerance was set at 20 ppm for the precursor ions and 0.8 Da for the ITMS fragment ions. Andromeda search included specific trypsin enzyme with maximum two missed cleavages, minimum of seven amino acids in length. Fixed modification carbamidomethyl (C), and variable modifications of oxidation (M), acetyl (Protein N-term), and deamidation (NQ) were considered. Protein identification threshold was set to 1% false discovery rate (FDR) as described previously (Cox and Mann, 2008).

Proteins that exhibit statistically significant abundance between CDH2-BioID2 to control were selected as follows. Proteins with a single peptide identification were excluded from the data analysis and Student’s t-test on log2 transformed protein intensity was used for the statistical inference to select CDH2-BioID2 interacting proteins. A protein was considered a significant candidate if the t-test p-value was <0.001 and the fold change >10 when compared to the control.

As a surrogate for protein abundance, MaxQuant iBAQ values were used for label-free absolute quantification of identified proteins (Schwanhausser et al., 2011). The average iBAQ value for each protein was determined from the six replicates in the both CDH2 and control samples. The final iBAQ value (provided in Table S1) was determined by subtracting the control average from the CDH2 average.

### Bioinformatics analysis

CDH1 BioID proximity proteomics results were from two previous studies (Guo et al., 2014; Van Itallie et al., 2014). The ICD protein list was from a previous curation (Estigoy et al., 2009). Venn diagrams comparing the protein lists were generated using BioVenn (Hulsen et al., 2008)(Hulsen et al., 2008)(Hulsen et al., 2008). Pathway and disease & function enrichment analysis was performed using Ingenuity Pathway Analysis (IPA) tools (Qaigen). Gene expression data for the identification of HEGs (Heart Enriched Genes) or CEGs (Cardiomyocyte Enriched Genes) were from previous studies (Fagerberg et al., 2014; Li et al., 2017; Li et al., 2015; Tompkins et al., 2016). The heart or cardiomyocyte enriched genes were identified by the Gene Expression Pattern Analyzer (GEPA) algorithm at the threshold of fold change ≥2 (≥2.5 for cardiovascular differentiation from human embryonic cell data set) (Li et al., 2015). Three types of expression patterns were selected: 1) exclusive high expression in cardiomyocytes or heart; 2) multiple high expression tissues/cells including heart/cardiomyocytes in which the sum of fragment per kilobase of exon per million reads (FPKM) was greater than the total sample number and the number of pattern samples was no greater than 4; and 3) “gradient” pattern with the highest expression in heart/cardiomyocytes and fold change of the highest and lowest expression is no less than 4. Fisher’s exact tests for overrepresentation analysis of HEGs or CEGs were performed using R.

### Protein Network Analysis

The protein interaction map was generated using IPA pathway designer. Only experimentally supported protein-protein interactions were considered. Hierarchical classification was done by grouping the proteins manually using CDH2 at the core. Proteins that bind CDH2 directly were designated as primary interactors. Proteins that bind to primary interactors but not CDH2 were classified as secondary interactors. Proteins that bind secondary interactors were designated as tertiary interactors. Finally, proteins that bind tertiary interactors or to outermost tier proteins were defined as quaternary interactors. 52 proteins could not be linked to the protein network.

## Acknowledgments

We are grateful to Tanya Svitkina for her guidance with the PREM experiments. We thank Ulf Schwarz and Kristofer Fertig (Leica Microsystems) for assistance with STED imaging.

## Competing Interests

No competing interest declared.

## Funding

This work was supported by National Institutes of Health F31 HL136069 (C.D.M.) and R01 HL127711 (A.V.K). STED imaging was supported by National Institutes of Health 1S10OD021540 (S.C.W.). This project used the Hillman Cancer Proteomics Facility / Biomedical Mass Spectrometry Center that is supported in part by National Cancer Institute P30CA047904.

## Supplemental Figures

**Supplemental Figure 1.**
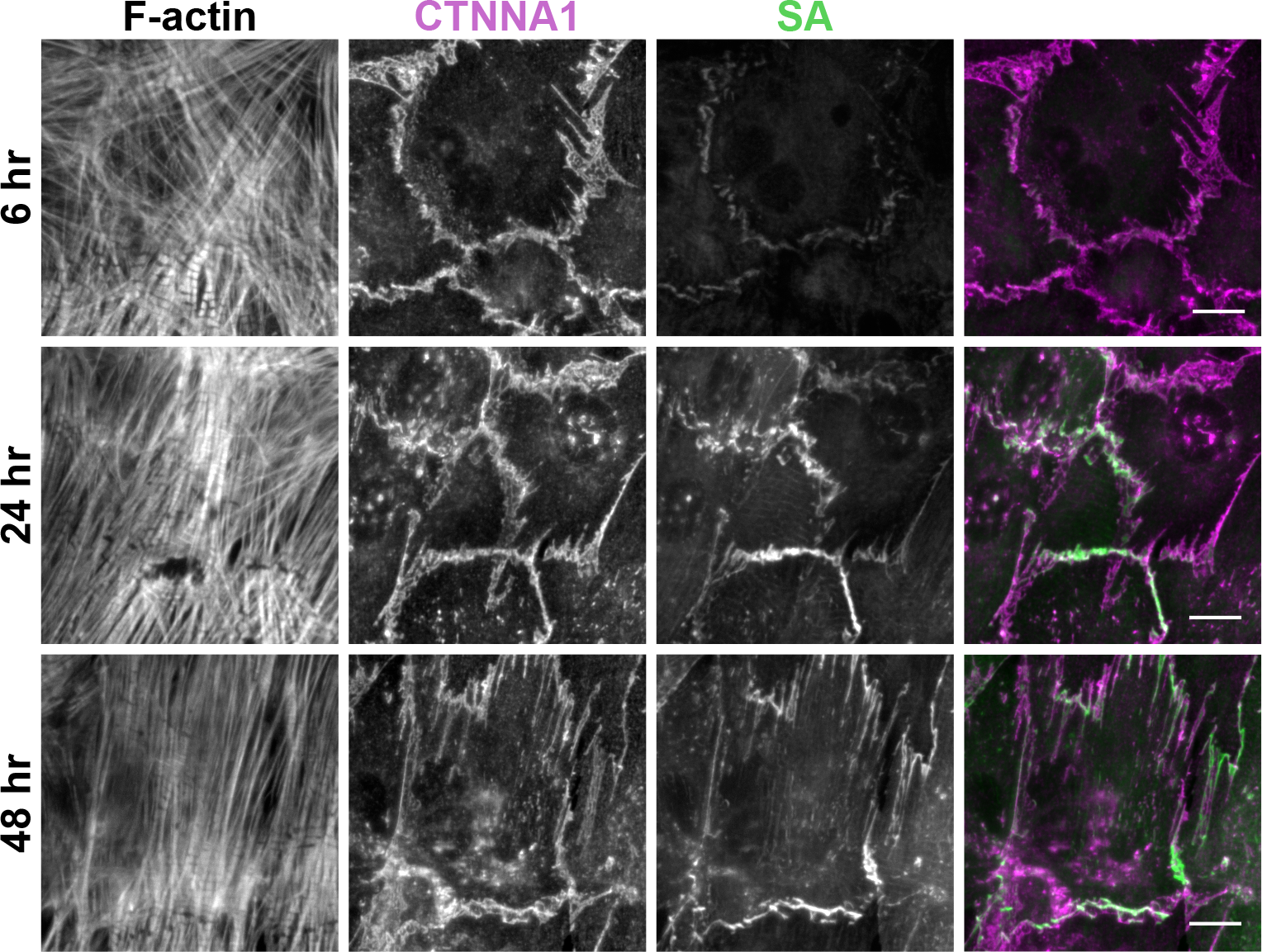
(accompanies Fig 3) Time course of biotin labeling. Cardiomyocytes infected with Cdh2-BioID2 adenovirus were fixed 6, 24 and 48 hours after biotin addition. Cells were stained for F-actin, CTNNA1 and biotin (streptavidin, SA). CTNNA1 (magenta) and SA (green) channels are shown in merge. Scale bar is 10 μm.

**Supplemental Figure 2.**
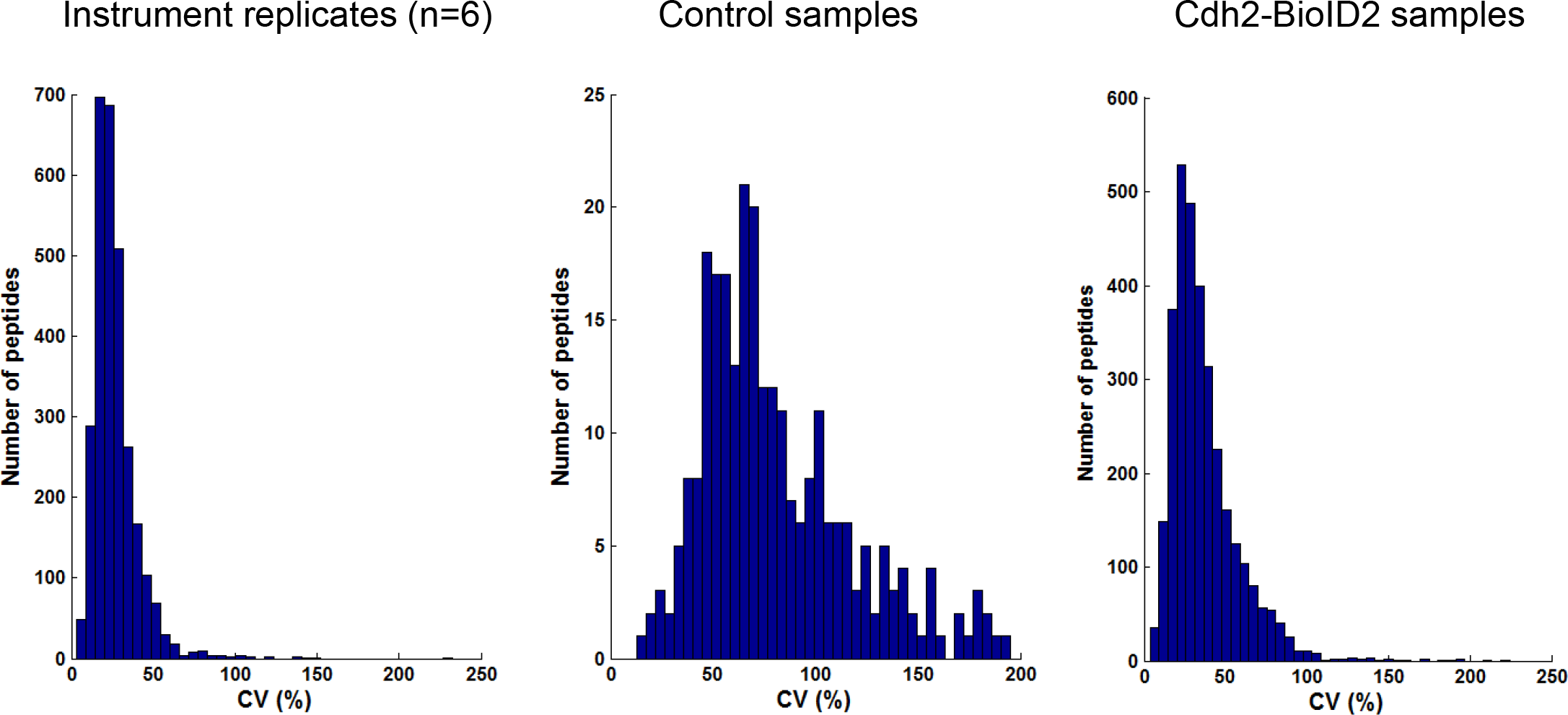
(accompanies Fig 4). Coefficient of variance (CV) for mass spec analysis of instrument replicates, control samples and experimental (Cdh2-BioID2) samples.

**Supplemental Figure 3.**
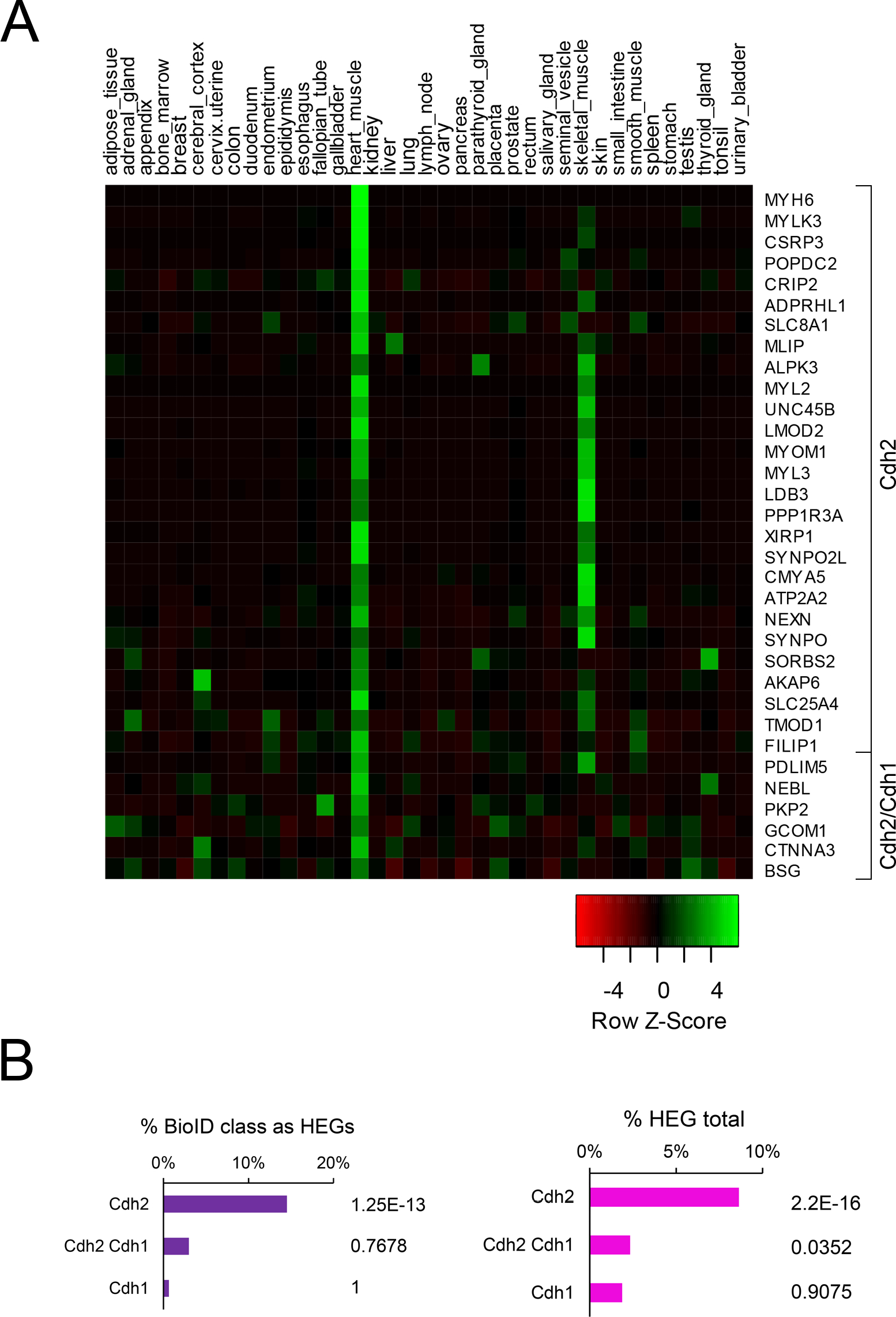
(accompanies Fig 5) A. Heat map of CDH2 or CDH2/CDH1 expression profiles in human tissues. B. Left, percentage of each BioID class as HEGs. Right, fraction of those BioID HEGs in the total HEG population. P value of Fisher’s exact test shown.

**Supplemental Figure 4.**
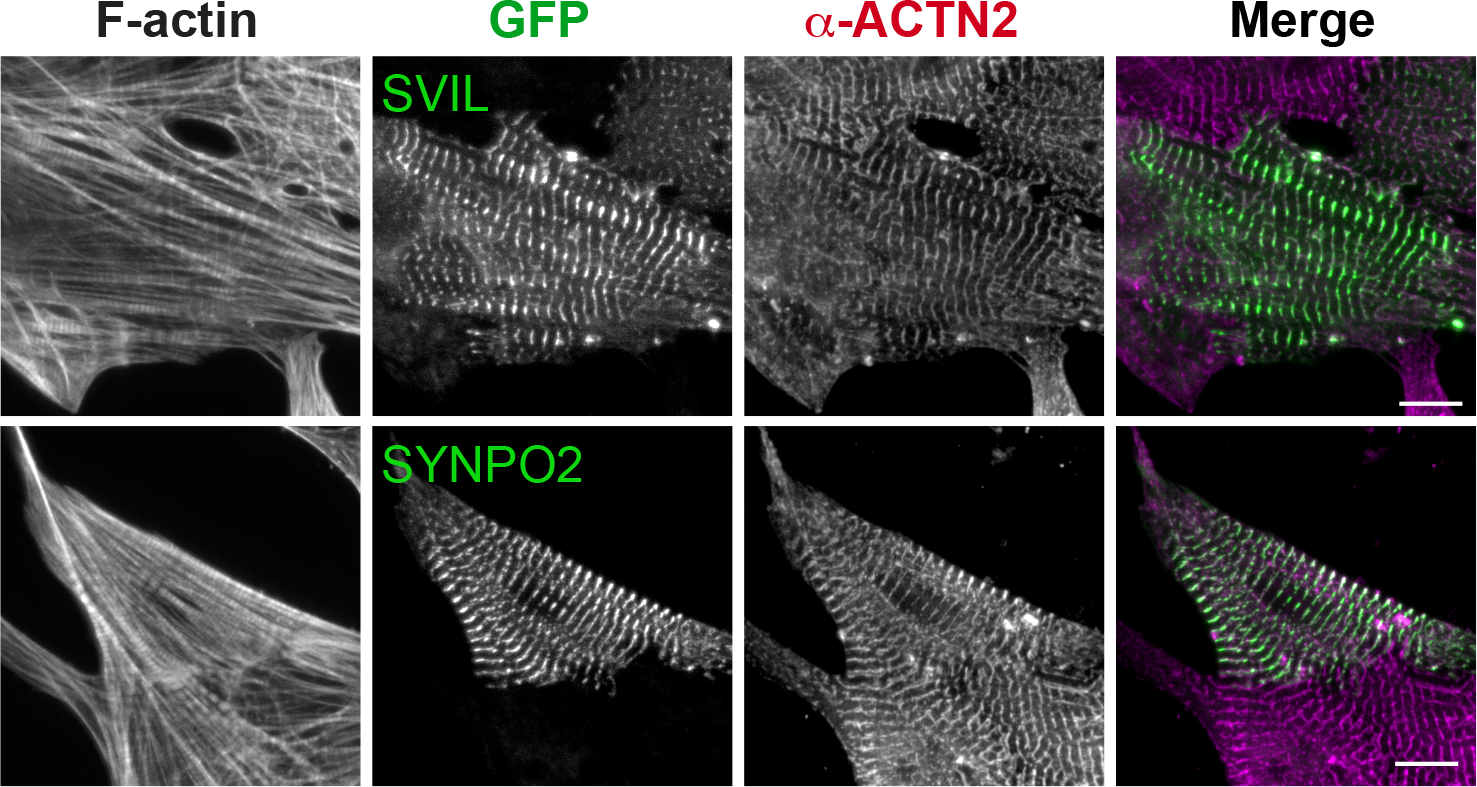
(accompanies Fig 7). SVIL and SYNPO2 localize to Z-discs. Cardiomyocytes transfected with EGFP-tagged SVIL and SYNPO2. Cells were fixed 24 hours post-transfection and stained for ACTN2 and F-actin. Scale bar is 10 μm.

**Supplemental Table 1**. Cardiomyocyte CDH2 Interactome.

**Supplemental Table 2**. IPA enrichment in canonical pathways

**Supplemental Table 3**. IPA enrichment in disease and function

**Supplemental Table 4**. Cdh2-BioID2 hits unconnected to the protein-protein interaction map

